# Transmission of Mixed Convergent Signals at the Mouse Retinogeniculate Synapse

**DOI:** 10.1101/2024.08.01.606255

**Authors:** Takuma Sonoda, Qiufen Jiang, Héctor Acarón Ledesma, Wei Wei, Chinfei Chen

**Author notes:** These authors contributed equally to this work (Takuma Sonoda, Qiufen Jiang).

## Abstract

There are two broad modes of information transfer in the brain: the labeled line model, where neurons relay inputs they receive, and the mixed tuning model, where neurons transform and integrate different inputs. In the visual pathway, information transfer between retinal ganglion cells (RGCs) and the dorsal lateral geniculate nucleus (dLGN) neurons is primarily viewed as a labeled line. However, recent work in mice has demonstrated that different RGC types, encoding distinct visual features, can converge onto a dLGN neuron, raising the fundamental question of whether the dLGN transforms visual information. Using optogenetics we activated distinct RGC populations and assessed spiking output of dLGN neurons by *in vivo* recordings. We found that visual response properties of dLGN neurons driven by a specific RGC population largely matched properties of the activated RGCs. Furthermore, *in vitro* dual-opsin experiments demonstrate that strong functional convergence from distinct RGC types rarely occurs. Thus, retinogeniculate information transfer in mice largely adheres to a labeled line model.

## Introduction

In the primary pathway mediating visual perception, retinal ganglion cells (RGCs) synapse onto thalamocortical (TC) neurons in the dorsal lateral geniculate nucleus (dLGN), which then transmit information to the primary visual cortex (V1) (Usrey and Alitto, 2015; Seabrook et al., 2017; Hooks and Chen, 2020). Classic functional studies in the cat visual system demonstrated that TC neurons receive strong "driver-like" inputs from a few RGCs that are capable of evoking spiking (Bishop et al., 1958; Hubel and Wiesel, 1961; Cleland et al., 1971; Mastronarde, 1992; Usrey et al., 1998, 1999). These findings have established the prevailing view that TC neurons function as simple relays of retinal input. However, several of these studies also noted the presence of weak connections, indicating that TC neurons can receive weak functional inputs from numerous RGCs (Hubel and Wiesel, 1961; Cleland et al., 1971; Mastronarde, 1992; Usrey et al., 1998, 1999). In fact, most connected RGC-TC neuron pairs do not exhibit a 1:1 relationship in terms of spiking, as would be predicted by a simple labeled line model (Cleland et al., 1971; Usrey et al., 1998, 1999). Therefore, these studies raised the intriguing possibility that signals from multiple RGCs are combined in the dLGN.

Recent work in the mouse visual system has reignited interest in the idea that TC neurons might function as integrators rather than simple relays of RGC input. Studies examining the RGC inputs onto dLGN neurons have shown that more than 10 RGCs can innervate a single dLGN neuron (Hammer et al., 2015; Morgan et al., 2016; Litvina and Chen, 2017) and that these RGCs can consist of different types (Rompani et al., 2017; Liang et al., 2018; Jiang et al., 2022). These studies have been interpreted by many as evidence that mouse visual processing is significantly different from that of higher mammals. Indeed, extensive work has revealed important species differences, including lamination patterns and the relative abundance of RGC types in mice compared to higher mammals (Nassi and Callaway, 2009; Liang and Chen, 2020; Kerschensteiner, 2022). However, there has been a fundamental difference in the approaches used to understand visual processing in different species. For example, seminal cat or primate studies largely focused on dLGN neuron spiking responses using *in vivo* recording approaches (Bishop et al., 1958; Hubel and Wiesel, 1961; Cleland et al., 1971; Sincich et al., 2007), whereas work in mice has taken advantage of genetic tools and other modern approaches with higher resolution, sensitivity and specificity. Therefore, conclusions drawn about different species could largely be a product of the levels of analyses used to study visual processing.

To narrow the gap between approaches used between species, we assessed the contribution of distinct RGC types to the spiking output of dLGN neurons in mice. We utilized optogenetics to activate subsets of RGCs *in vivo* and recorded resulting spiking activity in postsynaptic dLGN neurons. We found that the visual response properties of dLGN neurons that reliably spiked in response to optogenetic stimulation matched the properties of the RGC population being activated. Our findings suggest that despite the convergence of multiple RGC types onto a dLGN neuron, one type tends to dominate the postsynaptic spiking response. We also examined functional convergence of two distinct RGC types *in vitro* using a dual-opsin approach. When comparing synaptic strengths of the two distinct RGC types onto a given TC neuron, the likelihood of a cell receiving strong inputs from both populations is rare, consistent with a labeled line model. Our study suggests that despite species differences, the principles underlying information transfer in the retinogeniculate pathway remain largely conserved across species. Taken together with other published studies, our results raise future questions regarding the purpose of the weak convergent RGC inputs.

## Results

### Optogenetic stimulation of RGC axons in the dLGN in vivo

We first sought to examine connections between specific RGC populations and dLGN neurons by utilizing optogenetics *in vivo*. This approach involved measuring responses of the same dLGN neurons to 1) drifting sine-wave gratings and 2) optogenetic activation of a subset of RGC axons (Fig. 1a)—enabling us to functionally classify dLGN neurons that receive driving input from specific RGC populations. We validated this approach by making *in vivo* dLGN recordings in anesthetized *Chx10-Cre; ChR2* mice: a mouse line in which all RGCs express channelrhodopsin- 2 (ChR2) (Rowan and Cepko, 2004; Litvina and Chen, 2017) (Fig. 1a-b). Multisite silicone probes with an attached optical fiber (optoelectrodes) were used to make *in vivo* extracellular recordings from dLGN neurons while optogenetically activating RGC axons expressing ChR2 (Fig. 1a). Gratings were presented before and after a block of optogenetic trials to ensure that responses to optogenetic stimulation were not due to unstable recording conditions (Fig. 1c). Only units that exhibited reliable visual responses before and after optogenetic stimulation were included for analysis (see methods).

**Figure 1:**
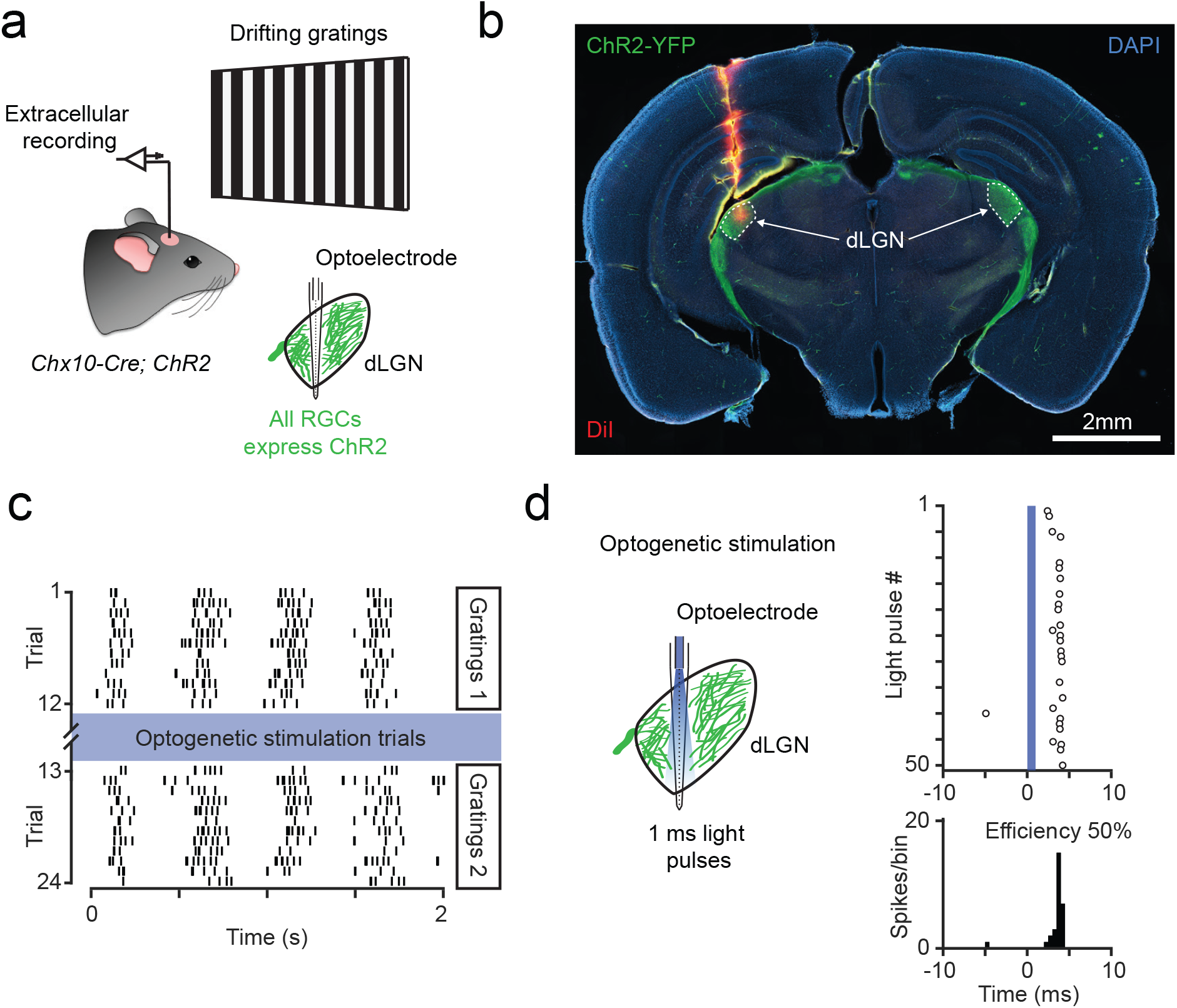
Measuring visual responses and functional inputs from RGCs simultaneously in dLGN neurons *in vivo*. **a,** *In vivo* extracellular dLGN recordings were performed in *Chx10-Cre; ChR2* mice in which all retinal cells, including RGCs, express ChR2. The axons of RGCs expressing ChR2 were optogenetically activated using silicon probes with optical fibers attached (optoelectrodes). This approach enabled simultaneous recording of dLGN neuron responses to visual stimuli (drifting gratings) and optogenetic activation of RGC axons *in vivo* **b,** Coronal brain section collected from a *Chx10-Cre; ChR2* mouse after *in vivo* dLGN recording. Green signals are RGC axons labeled in *Chx10-Cre; ChR2* mice. The electrode tract trajectory was labeled with DiI (red). This example shows the electrode clearly passing through the dLGN (dotted white border). **c,** Raster plot of responses to drifting gratings from an example dLGN neuron. Drifting gratings stimuli were presented before and after optogenetic stimulation trials (blue) to ensure recording stability. **d,** Responses of the same dLGN neuron in panel (c) to 1 ms optogenetic stimulation of RGC axons. The top raster plot shows spiking responses over a 20 ms window with time 0 indicating the onset of the light pulse. The bottom plot is a peristimulus time histogram of spiking activity in 0.5 ms bins. This neuron exhibited an opto-spike efficiency of 50% with a latency of 4 ms. Opto-spike efficiency was calculated by summing the number of spikes in the peak bin and the two adjacent bins, then dividing this sum by the total number of light pulses delivered (50 pulses). This value was then multiplied by 100 to convert it to a percentage.

During optogenetic stimulation trials, we activated RGC axons by delivering 1 ms light pulses at 5 Hz while animals viewed a dark screen. We quantified the extent to which spiking was evoked in postsynaptic dLGN neurons in response to RGC axon stimulation by binning spiking responses to optogenetic stimulation into 0.5 ms intervals to create a peristimulus time histogram (Fig. 1d). We defined opto-spike efficiency by summing the spikes in the peak interval with spikes in the two adjacent intervals, dividing this number by the total number of light pulses delivered, and multiplying by 100 to convert to a percentage.

On average, the first pulse of the 5 Hz pulse train elicited the most reliable spiking with an opto-spike efficiency of 37.3 ± 5.0% (mean ± SD, n = 37 visually responsive units) and delay of 4.4 ± 1.0 ms (Extended Data Fig. 1), consistent with a monosynaptic response (Usrey et al., 1998). No spiking was elicited in response to optical stimulation in Cre-negative *ChR2* animals (n = 7 units, Extended Data Fig. 1). These experiments demonstrate the feasibility of simultaneously measuring the visual response properties and the driving inputs of specific RGC types onto the same dLGN neuron *in vivo*.

### Optogenetic activation of on-off direction selective RGCs (ooDSGCs) elicits robust spiking in a subset of DS dLGN neurons

We next investigated the extent to which activation of a single RGC population can drive spiking in postsynaptic dLGN neurons *in vivo*. We utilized *Cart-IRES2-Cre* mice crossed to *Ai32* mice (*Cart-Cre; ChR2*) to express ChR2 in ooDSGCs. We have previously shown that this cross specifically labels ooDSGC inputs to the dLGN (Jiang et al., 2022). We broadly categorized the visual responses of dLGN neurons to drifting sine-wave gratings as broadly-tuned (BrT, 225/349 of visually responsive units), axis-selective (AS, 89/349 units), direction-selective (DS, 20/349 units), or suppressed by contrast (Sbc, 15/349) (Fig. 2a, c). The proportions of these functional categories were largely consistent with those reported by previous studies recording dLGN responses in similar conditions (Piscopo et al., 2013; Scholl et al., 2013; Zhao et al., 2013). DS dLGN neurons were defined as units exhibiting a direction-selectivity index (DSI) > 0.33; AS neurons were defined as units exhibiting a DSI < 0.33 and an axis-selectivity index (ASI) > 0.33 (Fig. 2c) (see methods).

**Figure 2:**
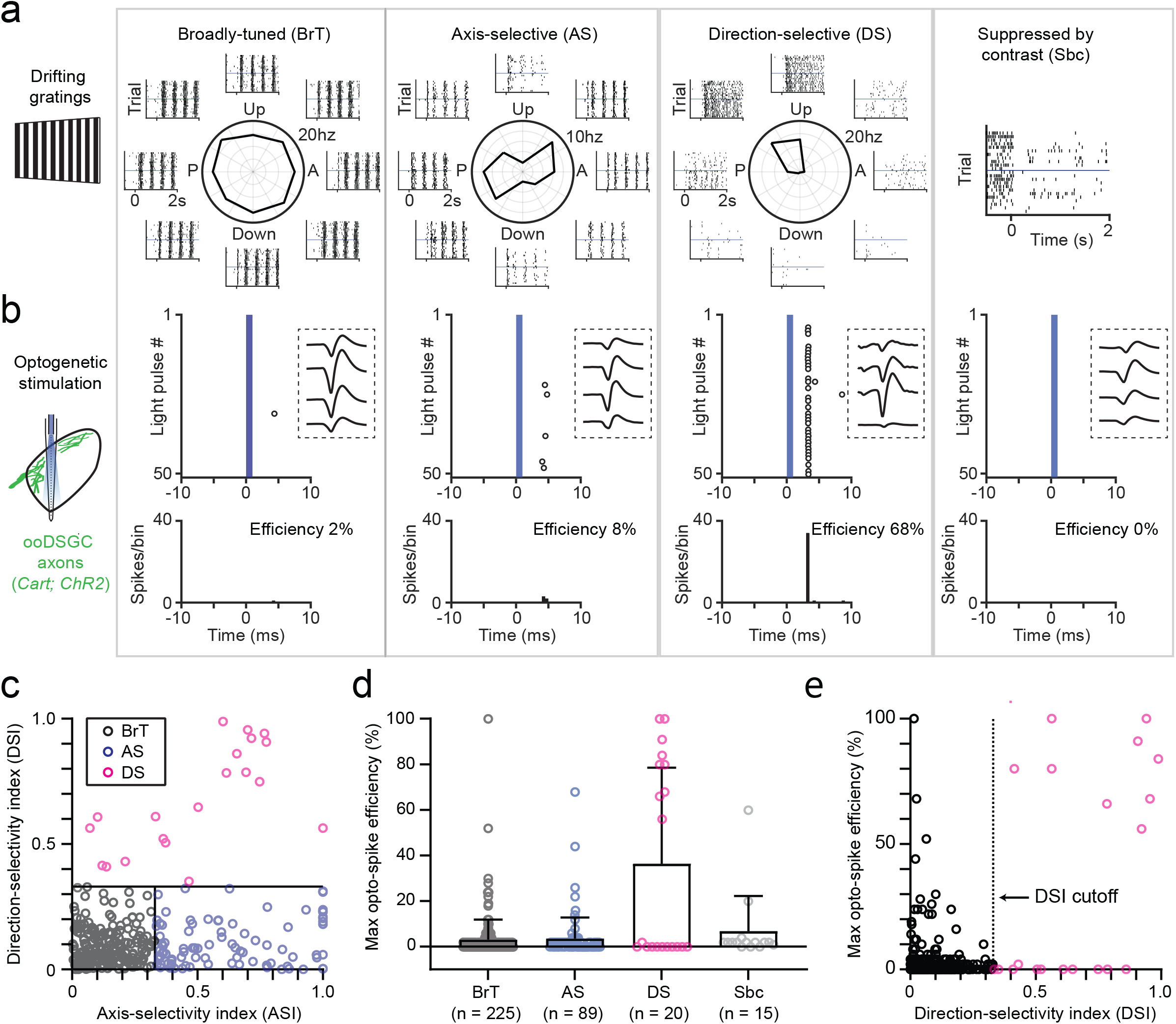
Visual responses of dLGN neurons receiving driving inputs from Cart-Cre^+^ RGCs. **a**, Example responses of broadly-tuned (BrT), axis-selective (AS), direction-selective (DS), and suppressed by contrast (Sbc) dLGN neurons to drifting gratings. Polar plots in the center represent average firing rate in response to gratings moving in 8 different directions. Raster plots surrounding each polar plot each show spiking activity in response to drifting gratings moving in a single direction across multiple trials. Time 0 denotes the onset of the drifting grating stimulus. Responses to gratings moving in a single direction is displayed for the Sbc dLGN neuron. **b,** Responses of the same neurons in panel (a) responding to optogenetic activation of Cart-Cre^+^ RGC inputs. The top raster plots show spiking responses in response to 1 ms light pulses over a 20 ms window. Open circles represent spikes. The onset of optogenetic stimulation occurs at time 0 and is represented by the solid blue box. The insets surrounded by dotted boxes are the average spike waveforms of each neuron recorded across multiple recording sites on the multielectrode array. The bottom plots are peristimulus time histograms of spiking activity in 0.5 ms bins. **c,** Scatter plot of ASI and DSI plotted for every visually responsive dLGN neuron with reliable spiking across trials, excluding Sbc neurons. BrT neurons (gray) were classified as cells with DSI < 0.33 and ASI < 0.33, AS neurons (blue) were classified as cells with ASI > 0.33 and DSI < 0.33, and DS (pink) neurons were classified as cells with DSI > 0.33. **d,** Max opto-spike efficiency of different functional classes in response to optogenetic activation of Cart-Cre^+^ RGC input. Data are from single units pooled from 38 *Cart-Cre; ChR2* animals. DS dLGN neurons are the primary population of dLGN neurons to receive driving input from Cart-Cre^+^ RGCs. **e,** Max opto-spike efficiency as a function of DSI. The dotted vertical line depicts the DSI cutoff used to classify DS neurons in this study (0.33).

Optogenetic activation of Cart-Cre^+^ axons in the dLGN elicited robust spiking, defined as an opto-spike efficiency greater than 40% (Fig. 2b, d), in only a small subset of dLGN units. We assessed which functionally defined group of dLGN neurons receive the strongest drive from ooDSGCs. On average, optogenetic activation of Cart-Cre^+^ RGC axons elicited the most reliable spiking in DS dLGN neurons compared to other functional types (Fig. 2a-d). 64% (9/14) of dLGN neurons that exhibited an opto-spike efficiency greater than 40% were DS (Fig. 2d). This specificity of Cart-Cre^+^ input for DS dLGN neurons did not depend on the DSI cutoff used for classifying DS dLGN neurons (0.33) (Fig. 2e). These results suggest a high degree of specificity in retinogeniculate connectivity when examined at the level of functional connections capable of driving spiking.

### Functional properties of DS neurons in the dLGN

Our results show that DS dLGN neurons exhibit a bimodal distribution along the parameter of opto-spike efficiency (Fig. 2d) raising the question of what characteristics differentiate DS neurons that receive strong versus weak ooDSGC input. In our experiments, we measured dLGN neuron responses to two sets of drifting grating-stimuli with different spatial frequencies: the first set of gratings being coarser (0.04 cycles/degree) and the other finer (0.16 cycles/degree). The temporal frequency of both sets of gratings was fixed at 2 cycles/second, therefore coarser gratings also moved faster than finer grating (50 degrees/second versus 12.5 degrees/second).

Interestingly, DS dLGN neurons could be subdivided by their preference for these two sets of gratings (Extended Data Fig. 2). DS dLGN neurons that preferred slower, finer gratings, which we term Type 1 DS, responded robustly to optogenetic activation of Cart-Cre^+^ inputs. Conversely, DS dLGN neurons that preferred faster, coarser gratings, termed Type 2 DS, did not respond to optogenetic activation of Cart-Cre^+^ input (Extended Data Fig. 2). Therefore, the preference of DS dLGN neurons for slower, finer gratings predicted the strength of Cart-Cre^+^ input received.

The spatial frequency and speed preference of DS dLGN neurons that are strongly driven by Cart-Cre^+^ RGCs is consistent with responses of ooDSGCs described previously (Wei, 2018; Summers and Feller, 2022). We directly confirmed this by imaging calcium responses of ooDSGCs in *Cart-Cre; GCaMP6f* mice to the same drifting gratings stimuli. ooDSGCs exhibited the same preference for slower, finer (0.16 cycles/degree) gratings—matching the preference of postsynaptic DS dLGN neurons that are strongly driven by optogenetic activation of Cart-Cre^+^ RGCs (Extended Data Fig. 3). These results further support the idea that ooDSGC responses are largely inherited by a subset of dLGN neurons.

#### Optogenetic activation of α-RGCs elicits reliable spiking in broadly-tuned dLGN neurons

Our results suggest that DS inputs to the dLGN exhibit a high degree of specificity. But does this specificity generalize to other RGC types? To address this, we utilized *Kcng4-Cre* mice, which primarily label α-RGCs, which are broadly-tuned (Duan et al., 2014). We targeted Kcng4-Cre^+^ RGCs by making eye injections of AAVs (AAV2/7m8-CAG-DIO-ChRger2-YFP) that drive Cre- dependent expression of a ChR variant, ChRger2 (Bedbrook et al., 2019), between P20-30 and *in vivo* recordings were performed between P60-120. We performed eye injections of AAVs instead of crossing to *ChR2* reporter mice because this cross results in labeling of dLGN neurons (data not shown). 94% (32/34) of dLGN neurons that were strongly driven by optogenetic activation of Kcng4-Cre^+^ inputs were broadly-tuned (Fig. 3a-b). Further analysis of broadly-tuned neurons that were strongly driven by Kcng4-Cre^+^ RGCs, as defined as a opto-spike efficiency of greater than 40%, showed clear ON or OFF responses to full-field luminance flashes (Fig. 3c). Only 1/21 of neurons that were responsive to full-field flashes had an ON-OFF response, as defined as responses with an on-off index between -0.33 and 0.33 (see methods) (Fig. 3c). Because α-RGCs consist of only ON and OFF types (Pang et al., 2003; Krieger et al., 2017), these results suggest that there is not strong functional convergence of ON and OFF α-RGCs inputs onto a single dLGN neuron and that information encoded by α-RGCs is also relayed to visual cortex. Thus, when examined at the level of inputs capable of driving spiking, the mouse retinogeniculate synapse exhibits a high degree of specificity like in higher mammals.

**Figure 3:**
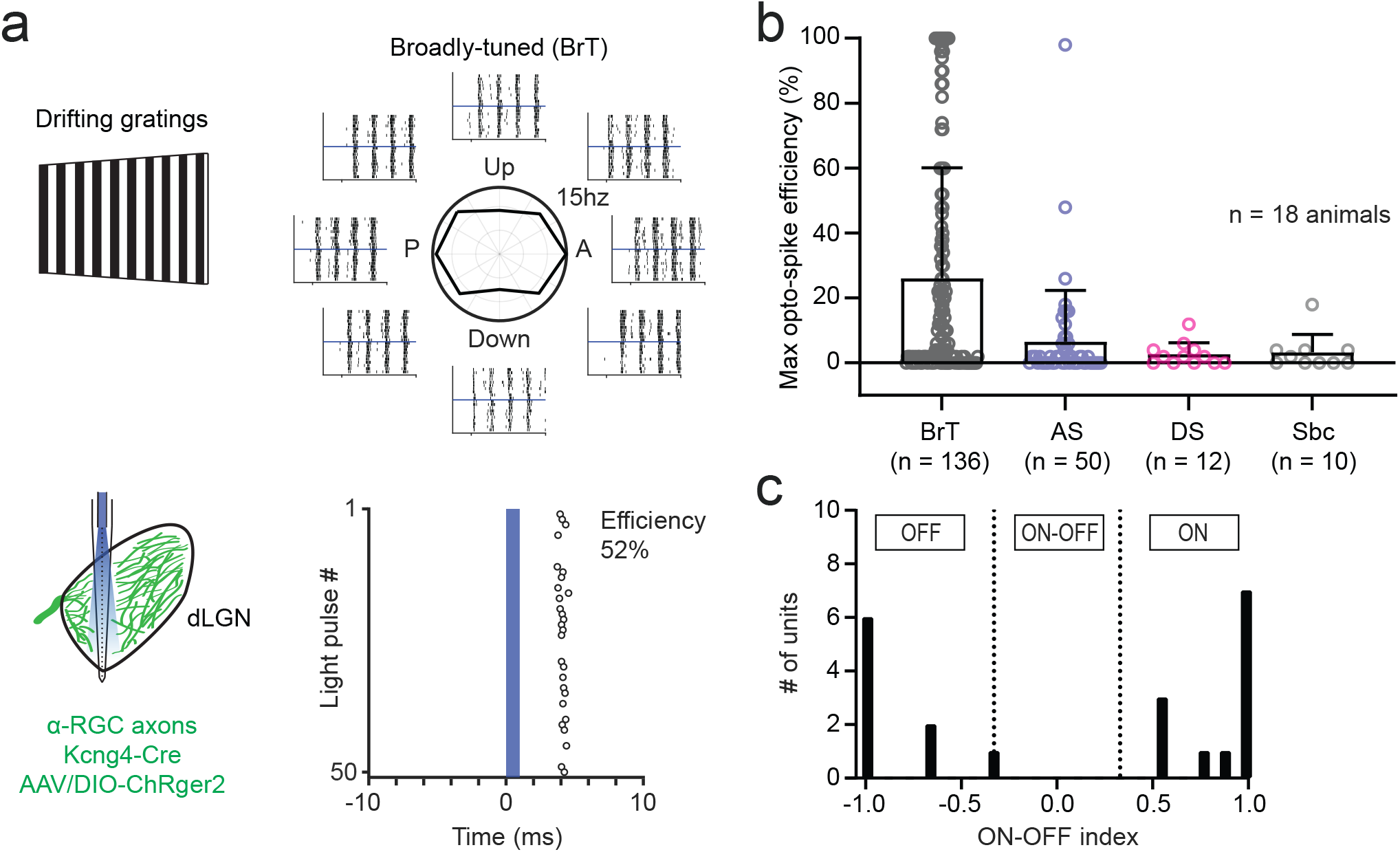
Visual responses of dLGN neurons receiving driving inputs from Kcng4-Cre^+^ RGCs. **a**, ChRger2, a ChR variant, was expressed in α-RGCs by injecting AAVs into *Kcng4-Cre* mice. The top plots are responses of a broadly-tuned dLGN neuron to drifting gratings moving in 8 directions. The bottom raster plot shows spiking responses of the same neuron in top panel to optogenetic activation of Kcng4-Cre^+^ RGC input over a 20 ms window. The onset of optogenetic stimulation occurs at time 0 and is represented by the solid blue box. **b,** Max opto-spike efficiency of different functional classes in response to optogenetic activation of Kcng4- Cre^+^ RGC input. BrT dLGN neurons are the primary population of dLGN neurons to receive driving input from Kcng-Cre^+^ RGCs. **c,** Histogram of on-off index measured in response to 1s full-field flashes. Data are from broadly-tuned dLGN neurons that received strong Kcng4-Cre^+^ inputs (opto-spike efficiency > 40%) (n = 21 units).

### Directly measuring functional convergence between α-RGCs and DS-RGC inputs

Our *in vivo* study shows that activating functionally defined populations of RGCs elicits reliable spiking in a small percentage of dLGN neurons, the majority of which exhibit the same functional properties as the optogenetically activated RGC population. These results suggest a simple inheritance of tuning features from the retina to the dLGN. However, it remains unclear whether dLGN neurons with clear visual preferences can still be driven by other information lines. To directly assess the potential of strong dominant inputs from different RGC inputs, we took advantage of a dual-opsin approach to label and activate two distinct RGC populations with different tuning features and measured their maximal synaptic drive.

We expressed a red-shifted opsin in α-RGCs by eye injection of *Kcng4-Cre* mice with AAVs that drive Cre-dependent expression of ChrimsonR-tdTomato (Klapoetke et al., 2014) (Fig. 4a). To label a RGC population distinct from α-RGCs, we co-injected AAVs that drive expression of a blue-shifted opsin, CatCh (Kleinlogel et al., 2011), in ooDSGCs under the control of the ProD1 promoter (Fig. 4a). ProD1 is a synthetic promoter that targets expression to a subset of ooDSGCs (Jüttner et al., 2019). Immunohistochemical analysis in the retina revealed that most of the CatCh-GFP^+^ RGCs (termed “ProD1^+^”) do not overlap with ChrimsonR-tdTomato^+^ RGCs (termed “Kcng4- Cre^+^”; 98% RGCs singly labeled) (Fig. 4b, c). Co-immunostaining for CART, which is a marker for ooDSGCs (Kay et al., 2011; Sabbah et al., 2017), shows that the majority (72%) of ProD1 labeled RGCs are CART-immunoreactive (CART-IR) and that 27% of CART-IR RGCs are ProD1^+^ (Fig. 4c-d). In the dLGN, the two different populations of RGCs exhibit distinct axon termination patterns in both coronal and parasagittal sections (Fig. 4e-f). ProD1^+^ labeled terminals localize to the superficial “shell” region of dLGN, while Kcng4-Cre^+^ axons terminate in a larger area spanning the “core” and “shell”, consistent with previous studies of ooDSGCs and α-RGCs (Cruz-Martín et al., 2014; Martersteck et al., 2017; Jiang et al., 2022) (Fig. 4e). To preserve the integrity of optic tract, we utilized parasagittal slices for voltage clamp recordings (Turner and Salt, 1998). In parasagittal slices, the terminal endings of ProD1^+^ and Kcng4-Cre^+^ RGCs intermingle in the ventral-posterior area (Fig. 4fi). Higher resolution images of this area reveal potential clusters of ProD1^+^ and Kcng4-Cre^+^ boutons (Fig. 4fii-iii), suggesting that these two different RGC types might converge onto the same dLGN neurons.

**Figure 4.**
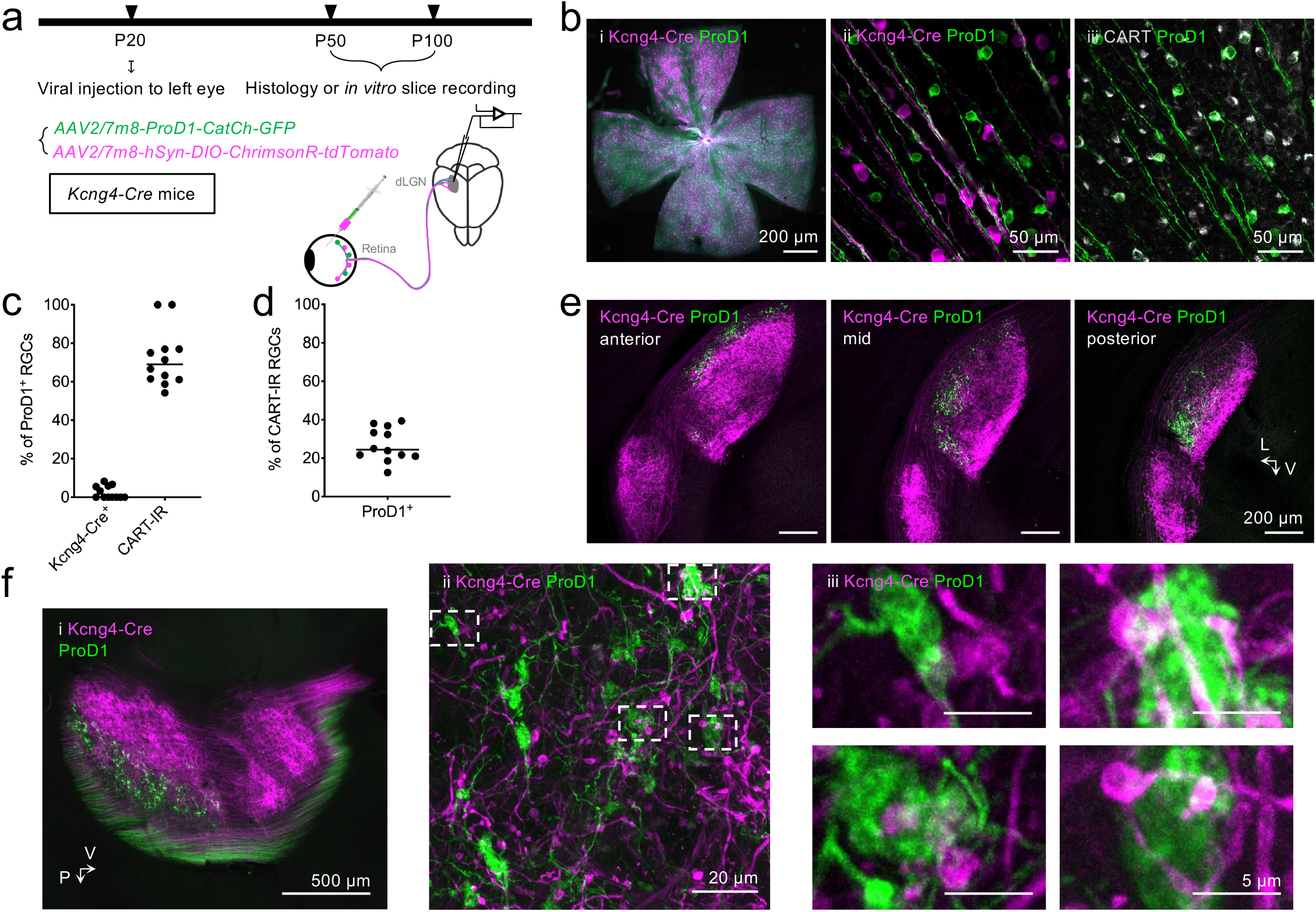
Viral strategy for simultaneously labeling !-RGCs and ooDSGCs in the retina. **a**, Strategy and experimental timeline for labeling Kcng4-Cre+ and ProD1+ RGCs in the retina. ∼P20 *Kcng4- Cre* mice were eye injected with viruses that drive Cre-dependent expression of ChrimsonR-tdTomato and Cre-independent *ProD1* promoter-driven expression of CatCh-GFP to label !-RGCs and ooDSGCs, respectively. Histology and *in vitro* slice recordings were performed between P50-100. **b,** Images of Kcng4-Cre+ and ProD1+ RGCs labeled in the retina. i) Low-magnification image of a whole-mount retina with Kcng4-Cre+ and ProD1+ RGCs labeled. Scale bar, 200 μm. ii) High-magnification image of the ganglion cell layer (GCL) of the retina showing ProD1+ (green) and Kcng4-Cre+ (magenta) RGCs exhibiting no overlap. Scale bar, 50 μm. iii) High-magnification image of the GCL showing a high degree of overlap between ProD1+ and CART-immunoreactive (CART-IR) RGCs (a marker of ooDSGCs). Scale bar, 50 μm. **c,** Proportion of ProD1+ RGCs that co-localize with Kcng4-Cre+ or CART-IR RGCs. The majority of ProD1+ RGCs (72%) are CART-IR and exhibit minimal overlap with Kcng4-Cre+ RGCs (2.5%). **d,** Proportion of CART-IR RGCs labeled by ProD1-promoter driven viral expression of CatCh-GFP. **e,** Coronal brain sections of the dLGN taken at different levels along the anterior-posterior axis. Kcng4-Cre+ (magenta) axons span both “shell” and “core” regions of the dLGN, while ProD1+ (green) axons are restricted to the “shell” region. L (lateral), V (ventral). Scale bar, 200 μm.

We next recorded from TC neurons in the ventral-posterior area of dLGN (Fig. 5a). We added Alexa Fluor 647 (Alexa647) in the internal solution to fill the neurons for posthoc morphological analysis. 3D reconstructions revealed the proximity of both Kcng4-Cre^+^ and ProD1^+^ axonal terminals to the proximal dendrites of post-synaptic dLGN neurons (Fig. 5a and Extended Data Fig. 5a). We measured the opto-stim responses of the two distinct RGC populations with different wavelengths of light (Fig. 5a). As CatCh can be activated by blue light (470 nm) and ChrimsonR is sensitive to both orange (> 600 nm) and blue light, synaptic responses from Kcng4- Cre^+^ inputs were evoked by a pulse of orange light (Fig. 5cii). Prolonged stimulation of orange light will drive ChrimsonR into the desensitized state, rendering it transiently unresponsive to blue light (Hooks et al., 2015). We took advantage of this property to obtain the response to ProD1^+^ inputs by delivering blue light stimulation immediately following 250 ms of orange light illumination (Fig. 5b-c). The amplitude of blue light-evoked currents is comparable to that of summed responses to Kcng4-Cre and ProD1 inputs obtained with a desensitizing orange light prepulse (Fig. 5d, *P* = 0.53, Wilcoxon matched-pairs signed rank test), suggesting that this approach is reliable in quantifying the synaptic strengths from the two distinct RGC populations. Comparison of Kcng4-Cre^+^ and ProD1^+^ input-evoked responses for each cell showed no significant difference in the averaged amplitude of optically evoked excitatory postsynaptic currents (oEPSCs) from the two populations (Fig. 5e, *P* = 0.57, Paired t-test). However, for most cells recorded, the oEPSC amplitudes are quite different for the two populations, with one population much greater than the other (Fig. 5e, f). Only a subset of TC neurons (35.2%) receive functional inputs from both Kcng4- Cre or ProD1 RGCs (Fig. 5f). However, many of these functional inputs were weaker than 600 pA, a current amplitude we have previously shown is needed to drive action potential firing of TC neurons (Liu and Chen, 2008). Only 5.56% of the TC neurons in the ventral-posterior area received inputs from both RGC populations that were greater than 600 pA (Fig. 5f). Taken together, these results indicate limited convergence between different information lines in the dLGN, favoring the labeled-line model in retinal transmission.

**Figure 5.**
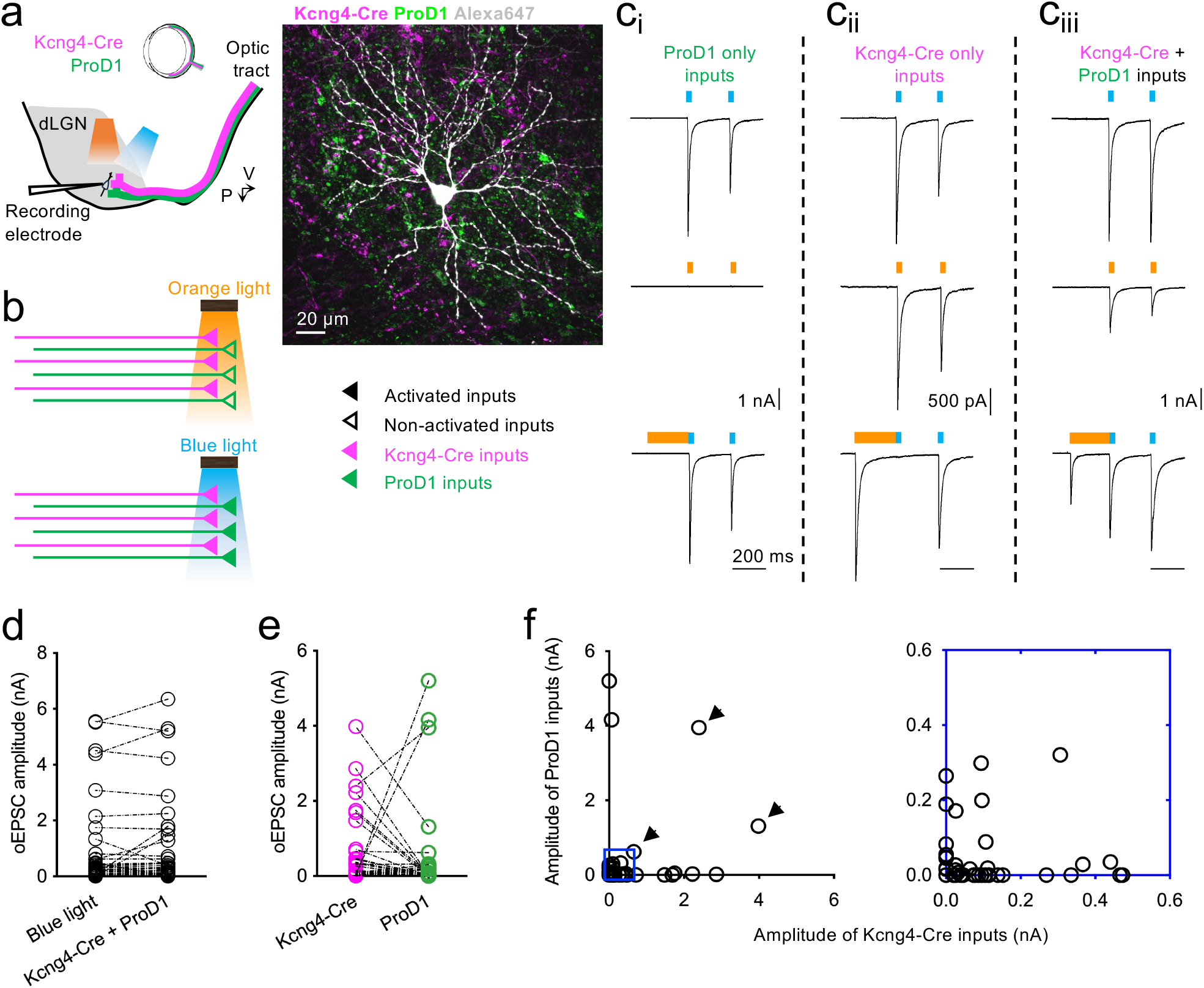
Skewed dLGN neuron responses to optogenetic activation of Kcng4-Cre^+^ and ProD1^+^ RGC inputs. **a,** Recording from TC neurons in ventral-posterior dLGN. *Left*: schematic showing recording configuration used for optical stimulation of Kcng4-Cre^+^ and ProD1^+^ inputs. Parasagittal sections of dLGN were prepared for patch clamp recording. Blue (470 nm) and orange (> 600 nm) full-field light stimulations were used for optogenetic activation. P (posterior), V (ventral). *Right*: Image showing an example TC neuron filled with Alexa Fluor 647 (Alexa647) to visualize dendritic morphology. Proximal dendrites were in close proximity to Kcng4-Cre^+^ and ProD1^+^ terminals. Scale bar, 20 μm. **b,** Schematic depicting activation of Kcng4-Cre^+^ and ProD1^+^ inputs with different wavelengths of light. Filled triangles represent activated terminals, while open triangles represent non-activated terminals. CatCh is expressed in ProD1^+^ axons (green) and can be activated by blue light (470 nm). ChrimsonR is expressed in Kcng4-Cre^+^ axons (magenta) and is activated by both orange (> 600 nm) and blue light. **c,** Examples of TC neurons that receive ProD1^+^ only input (i), Kcng4-Cre^+^ only input (ii), and both Kcng4- Cre^+^ and ProD1^+^ inputs (iii). To isolate the synaptic responses from the two populations of inputs, Kcng4- Cre^+^ inputs were evoked by a pulse of orange light (1 ms). Because blue light activates both inputs, a 250 ms of orange light illumination was first delivered to desensitize ChrimsonR in Kcng4-Cre^+^ axons rendering them transiently insensitive to optostim, followed by a 0.2 ms of blue light pulse. **d,** Amplitude of optically stimulated excitatory post-synaptic currents (oEPSCs) obtained by blue light stimulation compared to summed amplitude responses from isolated Kcng4-Cre^+^ and ProD1^+^ inputs. The two strategies for acquiring combined responses yield comparable amplitude of total oEPSCs. *P* = 0.53, Wilcoxon matched-pairs signed rank test. **f,** Axonal terminations of Kcng4-Cre+ and ProD1+ RGCs in a parasagittal dLGN section used for *in vitro* slice physiology recordings. i) Low-magnification image of the parasagittal dLGN with Kcng4-Cre+ and ProD1+ inputs labeled. Scale bar, 500 μm. ii) High-magnification overlay of Kcng4-Cre+ and ProD1+ boutons in ventral-posterior area of dLGN. Scale bar, 20 μm. iii) Potential clusters containing both Kcng4-Cre+ and ProD1+ boutons. Images were taken from areas indicated by dashed line boxes in panel (ii). Scale bar, 5 μm.

## Discussion

Previous studies using serial EM (Hammer et al., 2015; Morgan et al., 2016), single-rabies tracing (Rompani et al., 2017) and presynaptic imaging (Liang et al., 2018) suggested there is significant mixing of retinal signals in the mouse dLGN. Studies utilizing functional (Litvina and Chen, 2017) and modeling approaches (Rosón et al., 2019) showed that only 1-3 of these many RGC inputs onto a dLGN neuron are strong enough to influence spiking; however, it was still unclear what the nature of these strong inputs are. Do different RGC types drive spiking in a single dLGN neuron or do these strong inputs come from the same RGC type? In this study, we measured functional connections between specific population of RGCs and dLGN neurons using optogenetics *in vivo* and directly tested potential convergence of different RGC inputs *in vitro*. We focused on strong functional connections that elicit spiking, as has been the convention in functional studies of higher mammals. We report that at this level of analysis, the mouse retinogeniculate pathway is more consistent with a labeled-line model in which the dLGN relays retinal signals to V1. These findings should not diminish the importance of subthreshold inputs in visual processing but raise future questions regarding the purpose of weak convergent inputs.

### The mouse dLGN transmits information in a labeled line model

Our previous dLGN slice recordings in *Cart-Cre; ChR2* mice showed that while 79% of TC neurons receive functional Cart-Cre^+^ RGC input, only 38% of these cells receive strong inputs capable of driving spiking (Jiang et al., 2022). It was unclear whether these strong inputs dominate the postsynaptic response and result in DS being relayed to visual cortex. This was not possible to directly test in slice due to the lack of molecular or morphological markers to identify functional dLGN neuron types.

In this study, we use an *in vivo* approach to directly measure the functional output of dLGN neurons that receive strong inputs from Cart-Cre^+^ RGCs. If dLGN neurons receive strong inputs from different RGC types, we expected the visual responses of dLGN neurons driven by Cart-Cre^+^ RGCs inputs to be complex. However, we found that the primary population of dLGN neurons that are driven by Cart-Cre^+^ inputs match the functional properties of Cart-Cre^+^ RGCs (Fig. 2 and Extended Data Figs 2-3). These results demonstrate that RGCs of the same type likely provide strong inputs to dLGN neurons, which then relay these signals to visual cortex (Sun et al., 2016).

Interestingly, we observed a population of DS dLGN neurons that appear to lack a strong driver input from Cart-Cre^+^ RGCs (“Type 2 DS” in Extended Data Fig. 2). The fact that these Type 2 DS dLGN neurons do not receive input from Cart-Cre^+^ RGCs is consistent with their distinct stimulus preference for gratings that do not optimally activate ooDSGCs (Extended Data Fig. 2 and 3). These results raise the question of how DS is computed in Type 2 DS dLGN neurons. It is possible that DS is being driven by other DS RGC types, such as F-mini RGCs or ON-DSGCs; however, the speed tuning of F-mini and ON-DS RGCs suggests that this is unlikely (Rousso et al., 2016; Sivyer et al., 2019; Summers and Feller, 2022; Mani et al., 2023). Another possibility is that DS is computed locally via a disynaptic pathway involving interneurons. The existence of distinct populations of DS dLGN neurons indicates the dLGN may send multiple forms of DS signals to the visual cortex, as suggested by a previous study (Hillier et al., 2017).

We conducted experiments in *Kcng4-Cre* mice to determine whether our results in *Cart-Cre; ChR2* animals generalize to inputs from other RGC types. *Kcng4-Cre* mice primarily label α- RGCs, which are broadly-tuned. Importantly, we find that the primary population of dLGN neurons that are driven by activation of Kcng4-Cre^+^ inputs are also broadly-tuned (Fig. 3), further supporting the labeled line model for retinogeniculate processing to other RGC types. Moreover, Kcng4-Cre^+^ inputs primarily target the “core” region of the dLGN, while Cart-Cre^+^ inputs primarily innervate the superficial “shell” region (Fig. 4). This suggests that our results are applicable to both regions of the dLGN. As more genetic tools are developed to manipulate specific RGC types, it will be important to further explore how information encoded by other RGC types shapes visual responses of dLGN neurons.

Notably, a smaller fraction of dLGN neurons were driven by Cart-Cre^+^ RGCs using *in vivo* recordings (Fig. 2) compared to *in vitro* slice recordings (Jiang et al., 2022). These findings are consistent with the idea that the remaining fraction of Cart-Cre^+^ RGC are too weak to drive postsynaptic spiking. However, one caveat to this interpretation is that there are differences in dLGN regions sampling between the two methods. *In vivo* recordings span both “shell” and “core” regions of the dLGN (Fig. 1b), while slice recordings in Jiang et al. focused on the ventral-posterior region, which is enriched for Cart-Cre^+^ inputs. We do not believe this difference in dLGN regions can fully explain our results because our dual opsin experiments also showed infrequent strong inputs from two different RGC types despite spatial overlap (Fig. 5).

### Potential signal transformation in the dLGN

Our results generally support a labeled-line model for the transmission of retinal signals to visual cortex, but they leave room for the possibility of signal transformation within the dLGN. In our dual-opsin experiments, we focused recordings to the ventral-posterior region of the dLGN, which receives inputs from both Kcng4-Cre^+^ and ProD1^+^ RGCs (Fig. 4). In this region that is enriched for both inputs, 5.56% of neurons receive strong (> 600 pA) (Liu and Chen, 2008) input from both Kcng4-Cre^+^ and ProD1^+^ RGCs (Fig. 5), suggesting that a small percentage of dLGN neurons can integrate strong inputs from different RGC types. This may explain our *in vivo* data, where there were outliers in the BrT, AS, and Sbc dLGN populations that were strongly driven by Cart-Cre^+^ input (Fig. 2d). It is possible that these outliers are examples of dLGN neurons that integrate strong inputs from multiple RGC types. However, we have previously shown that the *Cart-IRES2- Cre* line labels ooDSGCs with an efficiency of 73% while also labeling 11.2% of BrT, 2.6% of AS and 9.3% of Sbc RGCs (Jiang et al., 2022). Therefore, we cannot rule out the possibility that these outliers are due to off-target labeling of RGCs.

Notably, this percentage measured functionally for convergent strong inputs is lower than the percentage predicted based on anatomical data (Hammer et al., 2015; Morgan et al., 2016; Rompani et al., 2017). This discrepancy between functional and anatomical data is further highlighted by an example recording where we performed a 3D reconstruction of a filled dLGN neuron with both Kcng4-Cre^+^ and ProD1^+^ terminals labeled (Extended Data Fig. 4a). Light microscopy analysis of spatially close contacts onto the dLGN neuron would predict comparable functional input from both RGC populations, however recording data from the same neuron revealed a strong input (812 pA) from only Kcng4-Cre^+^ RGCs (Extended Data Fig. 4b). Therefore, while anatomical data provide valuable information about potential connections, our study underscores the importance of understanding the strength of functional synaptic connections to fully understand information transfer in the brain.

Results from this study and previous studies (Litvina and Chen, 2017; Jiang et al., 2022) have shown that most RGC inputs are not capable of driving spiking, raising the question of what is the function of these abundant, subthreshold inputs? We previously showed that arousal and brain state can differentially alter the activity of axon terminals of specific RGC types in the dLGN (Liang et al., 2020; Reggiani et al., 2023). Specific RGC type modulation coupled with arousal-mediated changes in dLGN neuron excitability (Erisken et al., 2014) could result in inputs becoming more prominent in different contexts. *In vivo* recordings in this study, like those in cats, were conducted in anesthetized animals, limiting the influence of behavioral states on spiking output in our studies. An interesting avenue for future research will be to understand how specific connections are altered in different contexts.

Importantly, weak connections between RGCs and dLGN neurons are not a feature specific to the mouse retinogeniculate system, as these inputs have been described both at the functional and anatomical level in cats and primates (Cleland et al., 1971; Hamos et al., 1987; Usrey et al., 1998, 1999). In the cat visual system, these weak inputs have been hypothesized to be important sites of rapid plasticity that can be strengthened when a dLGN neuron loses its dominant RGC input (Moore et al., 2011). Further work is needed to understand the complex functions and computations that weak inputs may facilitate.

## Methods

### Animals

All procedures complied with the NIH Guide for the Care and Use of Laboratory Animals and were approved by the Institutional Animal Care and Use Committee (IACUC) at Boston Children’s Hospital. Both male and female mice were used, all on a C57 BL/6J background (Jackson Laboratories strain #000664). To express channelrhodopsin-2 (ChR2) in all RGCs, *Chx10-Cre* (JAX Strain #005105) (Rowan and Cepko, 2004; Litvina and Chen, 2017) mice were crossed to *Ai32* (JAX Strain #012569) (Madisen et al., 2012). To express ChR2 in on-off direction selective RGCs (ooDSGCs) for *in vivo* recording experiments, *Cart-IRES2-Cre-D* (JAX Strain #028533) were crossed to *Ai32* mice. For labeling and activating α-RGCs, *Kcng4-Cre* (JAX Strain #029414) (Duan et al., 2014) were used for both *in vivo* and dual-opsin slice recordings with intravitreal injection of AAVs (see below).

### Intravitreal injection of adeno-associate viruses (AAVs)

Intravitreal injections of AAVs were performed in *Kcng4-Cre* for both *in vivo* electrophysiology (Fig. 3) and dual-opsin *in vitro* slice experiments (Figs. 4-5). Mice were initially anesthetized in an isoflurane chamber (3.5% in oxygen) and isoflurane levels were typically maintained at ∼1.5% isoflurane in oxygen delivered through a nose cone. For *in vivo* recording experiments, the left eye was injected with 800 nL – 1 μL of AAV2/7m8-CAG-DIO-ChRger2-YFP (Addgene Plasmid #127329 packaged by BCH viral core). For dual-opsin experiments, the left eye was injected with 1 μL of a 1:1 mixture of AAV2/7m8-hSyn-DIO-ChrimsonR-tdTomato (Addgene Plasmid #62723 packaged by BCH viral core) and AAV2/7m8-ProD1-CatCh-GFP (Addgene Plasmid #125977 packaged by BCH viral core). Injections were made with a sharp glass pipette attached to either a 10 μL of Hamilton syringe or a Nanoject III microinjector (Drummond Scientific). Intravitreal injections were performed between postnatal day (P) 20-30. Following virus injection, *Kcng4-Cre* mice were housed under a standard 12:12 light dark cycle until experiments were performed between P50-100.

### Perfusion and immunohistochemical staining

For sampling of retinas or brain tissue to detect the expression of ChrimsonR-tdTomato and CatCh-GFP for dual opsin experiments, mice were euthanized with 10% pentobarbital and transcardially perfused with 0.1 M phosphate buffered saline (PBS) immediately followed by 4% w/v paraformaldehyde (PFA) in PBS. Eyes were removed and post-fixed in 4% PFA for 1 hr. Retinas were then dissected and rinsed in PBS before immunostaining. Brains were post-fixed overnight in 4% PFA at 4°C and rinsed in PBS. Brain slices containing dLGN were coronally sectioned through Leica VT1000 vibratome with thickness of 60 μm. Following *in vitro* electrophysiological experiments, parasagittal brain slices (250 μm) were collected, incubated in 4% PFA for 1 hr, and kept in PBS until immunostaining.

For whole-mount retina staining, retinas were blocked in PBS containing 5% normal goat serum (NGS) and 0.1% Triton X-100 at room temperature for 1 hr. Then primary antibodies were applied in PBS containing 0.3% Triton and 2% NGS: chicken anti-GFP (1:1000; ab13970, Abcam), rabbit anti-RFP (1:1000; 600-401-379, Rockland) or rabbit anti-CART (1:1000; H-003-62, Phoenix Pharmaceuticals), at 4°C for 2-3 days. After rinsing with 0.1% Triton/PBS, retinas were incubated with secondary antibodies at 4°C for another 2-3 days: goat anti-chicken antibody conjugated to Alexa Fluor 488 (1:1000; A11039, Invitrogen), goat anti-rabbit 555 (1:1000; A32732, Invitrogen) or goat anti-rabbit 647 (1:1000; A21245, Invitrogen). Retinas were then mounted and cover- slipped with Vectashield (H-1000, VectorLabs).

Coronal brain slices containing dLGN were stained with GFP and RFP overnight to label the axonal terminals from the dual opsins-labeled RGCs, followed by secondary antibody incubation for 2 hrs at room temperature. In electrophysiological experiments, patched cells were filled with Alexa Fluor 647 Hydrazide (1 mg/mL; A20502, Thermo Fisher). These parasagittal slices after patching were collected, stained with GFP and RFP following similar procedures to coronal brain slice staining, except that the incubation of primary and secondary antibodies was conducted at 4°C for 2-3 days each.

### Confocal microscopy

Whole-mount retina images were taken using a Zeiss Axio Imager.Z2 epifluorescence microscope. Z-stack and tile scan imaging of the whole-dLGN were performed using a Zeiss LSM980 Confocal with a 10x objective. To detect the expression of dual opsins in the retina of injected mice, images of the retina (12 fields of view from each mouse) were acquired through Zeiss LSM 710 Multiphoton Confocal using a 20x Olympus objective to detect GFP, RFP or CART signals. Quantification was performed manually using ImageJ. To examine potential convergence of Kcng4-Cre^+^ and ProD1^+^ boutons, and their interaction with filled post-synaptic neuron in the dLGN, higher resolution Z-stacks were taken under 63x objective. The cell and the retinal terminals were then re-constructed using Imaris 10.1.1 software.

### In vivo electrophysiology

Surgeries were performed in animals anesthetized with isoflurane (4 % for induction and 2 % for maintenance). Previous studies have shown that anesthesia does not substantially alter orientation- and direction-selectivity in the mouse dLGN, therefore we opted to perform recordings in anesthetized animals for recording stability (Zhao et al., 2013; Durand et al., 2016). 2.5 mg/kg dexamethasone was injected subcutaneously to prevent edema and eyes were lubricated with silicone oil to prevent drying. A 2-3 mm diameter craniotomy was performed and was centered

∼1.5 mm anterior and ∼2.5 mm lateral from the lambda suture. The exposed brain was always covered in artificial cerebral spinal fluid (aCSF) to prevent drying. The dLGN was targeted using stereotaxic coordinates 0.7 - 1.4 mm anterior and 2.10 - 2.45 mm lateral from the lambda suture. Up to 2 penetrations were made per animal and penetrations were spaced at least 0.2 mm apart in cartesian coordinates. Recordings were made from the right dLGN and the ipsilateral eye was covered during recordings.

32-channel silicon probes with an attached optical fiber were advanced slowly (<1 µm/second) until the entire dorsal-ventral limits of dLGN were included in within the boundaries of the electrode array. Two probe configurations were used in this study: 1) probes with recording sites arranged in a linear configuration with an attached 50 µm diameter optical fiber (Neuronexus, A1x32-5mm-25-177-OA32LP) and 2) probes with recording sites arranged in a staggered configuration with an attached 0.9 mm tapered fiber (Cambridge Neurotech, ASSY-37-H7b). Data were acquired using a PZ5 digitizer with RZ5P processor with Synapse software running on a Workstation 8 computer (Tucker-Davis Technologies). Signals were sampled at 24414 Hz and filtered between 300 Hz and 5 kHz to isolate single units.

### Visual stimuli and optogenetic stimulation in vivo

Visual stimuli were displayed on a 21 x 12 inch LED monitor, covering approximately 100° x 70° of the visual field. The screen was positioned 22 cm away from the animal’s left eye offset, at a 45-degree angle relative to the mouse’s anteroposterior axis. The screen had a mean luminance of 20 lux. Visual stimuli were gamma corrected and generated using Psychophysics toolbox (Brainard, 1997) in MATLAB (MathWorks).

To classify AS and DS dLGN neurons, full-field drifting sine-wave gratings moving in 8 directions were presented. These gratings consisted of 2 spatial frequencies (0.04 and 0.16 cycles/degree) with a temporal frequency of 2 Hz. Gratings were presented for 2 seconds with an interstimulus interval of 0.5 seconds. Unique spatial frequency and direction combinations were presented in pseudorandom order over 12 trials and a blank gray screen condition was included to measure spontaneous activity. Following these trials, a second set of gratings with identical parameters was presented after optogenetic stimulation. 20 trials of 1 second full-field luminance flashes were presented to measure ON-OFF responses.

*In vivo* optogenetic stimulation was delivered using a fiber-coupled 473 nm DPSS laser (Laserglow). During optogenetic stimulation trials, animals viewed a dark screen while 5 Hz pulse trains (each pulse lasting 1 ms) were delivered to activate RGC axons in the dLGN. 50 pulse trains were delivered during each recording session. To minimize confounding effects of ChR2 desensitization (Hooks et al., 2015; Litvina and Chen, 2017), there was a 10 second interval between each pulse train.

### Analysis of in vivo electrophysiology data

Spike sorting was performed using default parameters in Kilosort2 (Pachitariu et al., 2023). Kilosort outputs were manually curated in Phy (github.com/kwikteam/phy). Single unit responses to gratings were quantified by measuring the mean firing rate (F0) in response to all unique combinations of spatial frequency and direction. To determine if units were visually responsive, responses to blank gray screen trials were compared with responses from the spatial frequency/direction combination that elicited the largest response. A Mann-Whitney test was performed on responses to blank trials and preferred trials and units were classified as visual if *p* < 0.01. The preferred spatial frequency was determined by first measuring the direction that elicited the largest response. The spatial frequency that elicited the largest responses at the preferred direction was used as the preferred spatial frequency for each unit (Durand et al., 2016). To determine response reliability, a quality index (Qi) was computed as previously described (Baden et al., 2016). Spiking responses were segmented into 0.1 second bins to form a Trial x Response matrix, representing responses to the optimal frequency/direction gratings for each unit over multiple trials. Qi was calculated by dividing the variance of the mean response by the mean variance across trials. Units with a Qi less than 0.25 were excluded.

To assess unit stability before and after optogenetic stimulation trials, F0 responses from the preferred spatial frequency/direction combination were compared using a Mann-Whitney test. Units were excluded if F0 responses significantly differed (*p* < 0.05) after optogenetic stimulation trials. Additionally, Qi was measured by combining trials before and after optogenetic stimulation trials. Therefore, units that had significantly different responses after optogenetic stimulation were also excluded based on Qi (see above).

Axis-selectivity index (ASI) was computed as (Rpreferred – Rorthogonal) / (Rpreferred + Rorthogonal), where Rpreferred is the mean response to gratings moving along the cell’s preferred axis. Rorthogonal is the mean response to gratings moving along the axis orthogonal to Rpreferred. Direction-selectivity index (DSI) was computed as (Rpreferred – Ropposite) / (Rpreferred + Ropposite). Rpreferred is the mean response to gratings moving in the cell’s preferred direction (i.e. the direction that elicits the largest response). Ropposite is the response to gratings moving in the opposite of Rpreferred. Units with an ASI > 0.33 and DS < 0.33 were classified as AS; units with DSI > 0.33 were classified as DS; and units with DSI < 0.33 and ASI < 0.33 were classified as broadly-tuned. Units that exhibited responses that were significantly reduced by gratings were classified as Sbc.

ON-OFF index of units in response to full-field flashes was computed as follows: (RON - ROFF) / (RON + ROFF). RON and ROFF represent the mean firing rate of each unit calculated during the first 400ms following light onset (RON) and offset (ROFF). Units with an ON-OFF index greater than 0.33 were classified as ON; units with an ON-OFF index less than -0.33 were classified as OFF; and units with an ON-OFF index between -0.33 and 0.33 were classified as ON-OFF.

To quantify responses to optogenetic activation of RGC axons, we binned spiking responses to optogenetic stimulation into 0.5 ms intervals to create a peristimulus time histogram. The interval with the highest number of spikes within 10 ms of the optogenetic stimulus onset was used to measure synaptic delay. To calculate opto-spike efficiency, we summed the number of spikes in the peak interval with those in the adjacent intervals, divided this sum by the total number of light pulses (50), and multiplied by 100 to convert it to a percentage. This analysis was performed for each pulse in the 5 Hz pulse train, and the pulse with the highest opto-spike efficiency was used to determine the opto-spike efficiency for each unit. Typically, the first pulse in the 5 Hz pulse train elicited the most reliable spiking (Extended Data Fig. 1).

### Two-photon calcium imaging of ooDSGCs

Analysis of ooDSGC responses to drifting gratings was performed using our previously published dataset characterizing RGC types labeled in *Cart-IRES2-Cre* mice (Jiang et al., 2022). We analyzed a subset of ooDSGCs from this dataset that were presented with both moving bars and drifting grating stimuli matching those used for *in vivo* dLGN recordings.

### In vitro slice electrophysiology

Brain slices containing the optic tract (OT) and dLGN were prepared as previously described (Litvina and Chen, 2017; Jiang et al., 2022). Briefly, mice were anesthetized using isoflurane and decapitated into oxygenated (95% O2; 5% CO2) ice-cold cutting solution (in mM): 130 K-gluconate, 15 KCl, 0.05 EGTA, 20 HEPES, and 25 glucose (pH 7.4 adjusted with NaOH, 310-315 mOsm) (Pressler and Regehr, 2013). The brain was then removed quickly and immersed in the ice-cold cutting solution for 60 seconds. To obtain slices maintaining continuity of retinogeniculate fiber inputs, parasagittal sections were cut as previously described (Turner and Salt, 1998; Chen and Regehr, 2000). The brain was cut with a steel razor blade, then sectioned into 250 μm-thick slices in oxygenated, ice-cold cutting solution using a sapphire blade (Delaware Diamond Knives) on a vibratome (VT1200S; Leica). Slices containing the dLGN and OT were allowed to recover at 30°C for 15 minutes in oxygenated saline solution (in mM): 125 NaCl, 26 NaHCO3, 1.25 NaH2PO4, 2.5 KCl, 1.0 MgCl2, 2.0 CaCl2, and 25 glucose (pH 7.4, 310-315 mOsm).

To examine potential convergence of Kcng4-Cre^+^ and ProD1^+^ inputs in the dLGN, TC neurons from the ventral-posterior region were sampled for whole-cell patch clamp recording. Recordings were performed using a MultiClamp 700B (Axon Instruments), filtered at 1 kHz, and digitized at 20-50 kHz with an ITC-18 interface (Instrutech). Glass pipettes (Drummond Scientific) were pulled on Sutter P-97 Flaming/Brown micropipette puller (Sutter Instruments) and filled with internal solution containing (in mM): 35 CsF, 100 CsCl, 10 EGTA, 10 HEPES, and L-type calcium channel antagonist 0.1 methoxyverapamil (pH7.3, 290-300 mOsm) to optimize the pipette resistance to be 1.5-2.0 MOhm. To isolate excitatory synaptic currents, cells were recorded at room temperature in saline solution containing 20 μM of bicuculline (GABAAR antagonist; 0130, Tocris), 2 μM of CGP55845 (GABABR blocker; 1248, Tocris), 10 μM of DPCPX (antagonist of A1 adenosine receptors; 0439, Tocris), and 50 μM of LY341495 (blocker of presynaptic group II mGluRs; 1209, Tocris) (Kingston et al., 1998; Chen and Regehr, 2000; Hauser et al., 2013, 2014; Yang et al., 2014), to block inhibitory circuits and presynaptic neurotransmitter receptors that modulate retinogeniculate release (Chen and Regehr, 2000, 2003). Recordings were excluded if access resistance was larger than 10 MOhm or if access resistance changed by more than 20% during a recording.

### Dual opsin stimulation

To obtain the responses to two distinct RGC populations (Kcng4-Cre^+^ and ProD1^+^ RGCs) expressing different opsins, ChrimsonR and CatCh, different wavelengths of light were applied to evoke the excitatory post-synaptic currents (oEPSCs) in TC neurons. As CatCh can be activated by blue light (470 nm) and ChrimsonR is sensitive to both orange (> 600 nm) and blue light, synaptic response to ChrimsonR was evoked by a single pulse of orange light (1 ms) with full-field illumination through the 60x objective (Olympus LUMplanFL N 60x/1.00W). We found that prolonged stimulation of orange light drives ChrimsonR into the desensitized state and causes depolarization block of the pre-synaptic terminals, rendering it transiently unresponsive to blue light (Hooks et al., 2015). Therefore, the response to CatCh was obtained by delivering blue light stimulation (0.2 ms) immediately following 250 ms of orange light illumination. The blue light (83 mW) and orange light (102.4 mW) were supplied by a CoolLED pE-300^ultra^ unit. Stimulation duration ranged from 0.2 - 250 ms at highest power to obtain the maximal oEPSC.

### Analysis of slice electrophysiology data

Electrophysiological data acquisition and offline analysis were performed using custom software in IgorPro (Wave-Metrics). oEPSC amplitudes were obtained from average traces of 3-5 trials. Data calculation and statistical analysis were conducted using Graphphad Prism. All data sets were evaluated for normality using the Kolmogorov-Smirnov test. For nonparametric distributions, the Wilcoxon signed rank test were used for paired comparison. For normally distributed data sets, the paired t-test was used. For all figures, \**p* < 0.05; \*\**p* < 0.01; \*\*\**p* < 0.001.

---

**Extended Data Figure 1.**
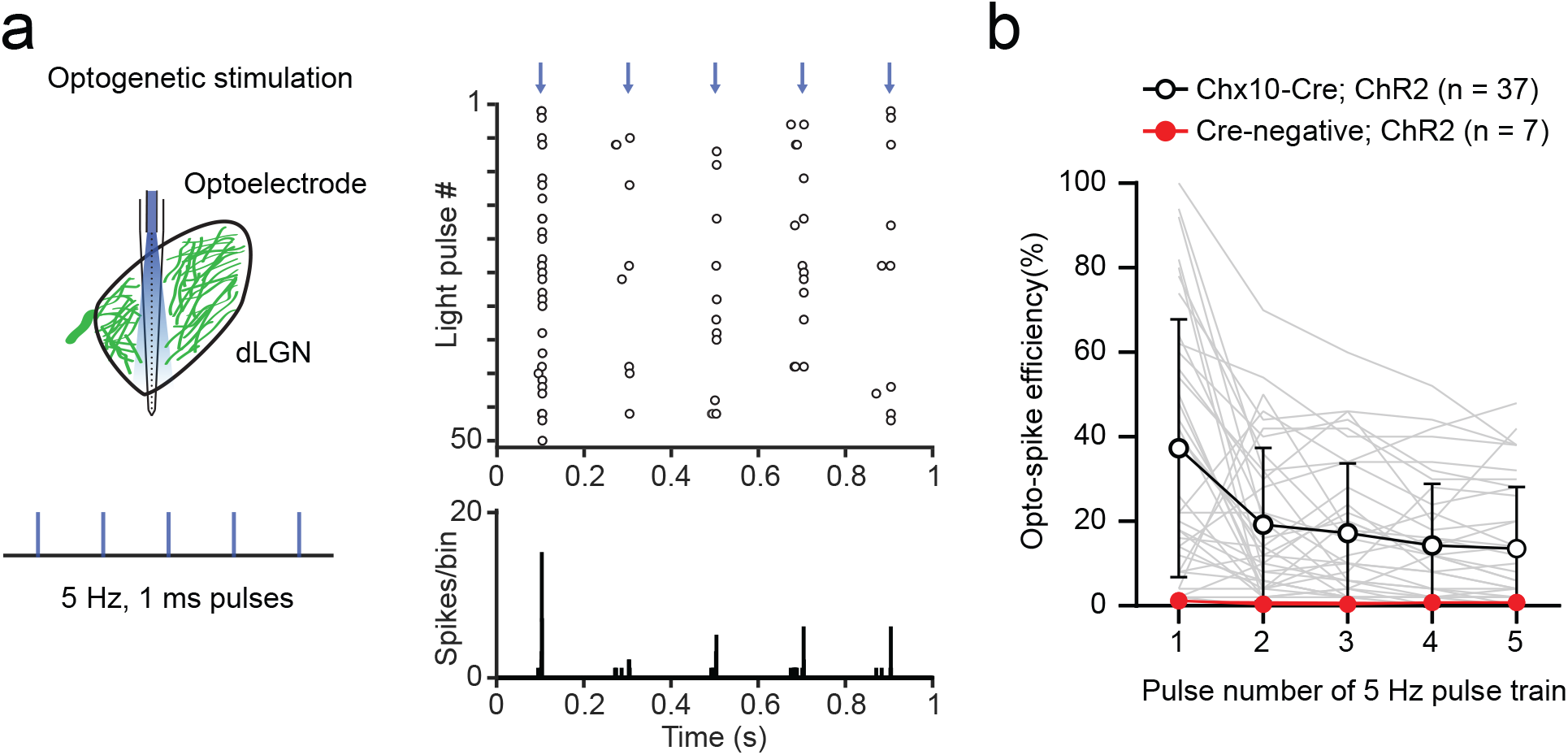
Responses of dLGN neurons to 5 Hz optogenetic stimulation in *Chx10-Cre; ChR2* mice. **a,** 5 Hz optical stimulation was used to activate RGC inputs during *in vivo* dLGN recordings in *Chx10; ChR2* mice. *Left*: Schematic of protocol used for 5 Hz train of optogenetic stimulation. *Right*: Raster plot with example responses of a dLGN neuron to 5 Hz stimulation across 50 trials. Blue arrows indicate when light pulses were delivered, and open circles represent spiking activity. The bottom graph shows peristimulus time histograms of spiking in 0.5 ms bins. **b,** Average opto-spike efficiency in response to each pulse in a 5 Hz pulse train. Gray lines represent responses of individual units to 5 Hz optical stimulation. Optical stimulation did not elicit robust spiking in control animals that did not express ChR2 (red).

**Extended Data Figure 2.**
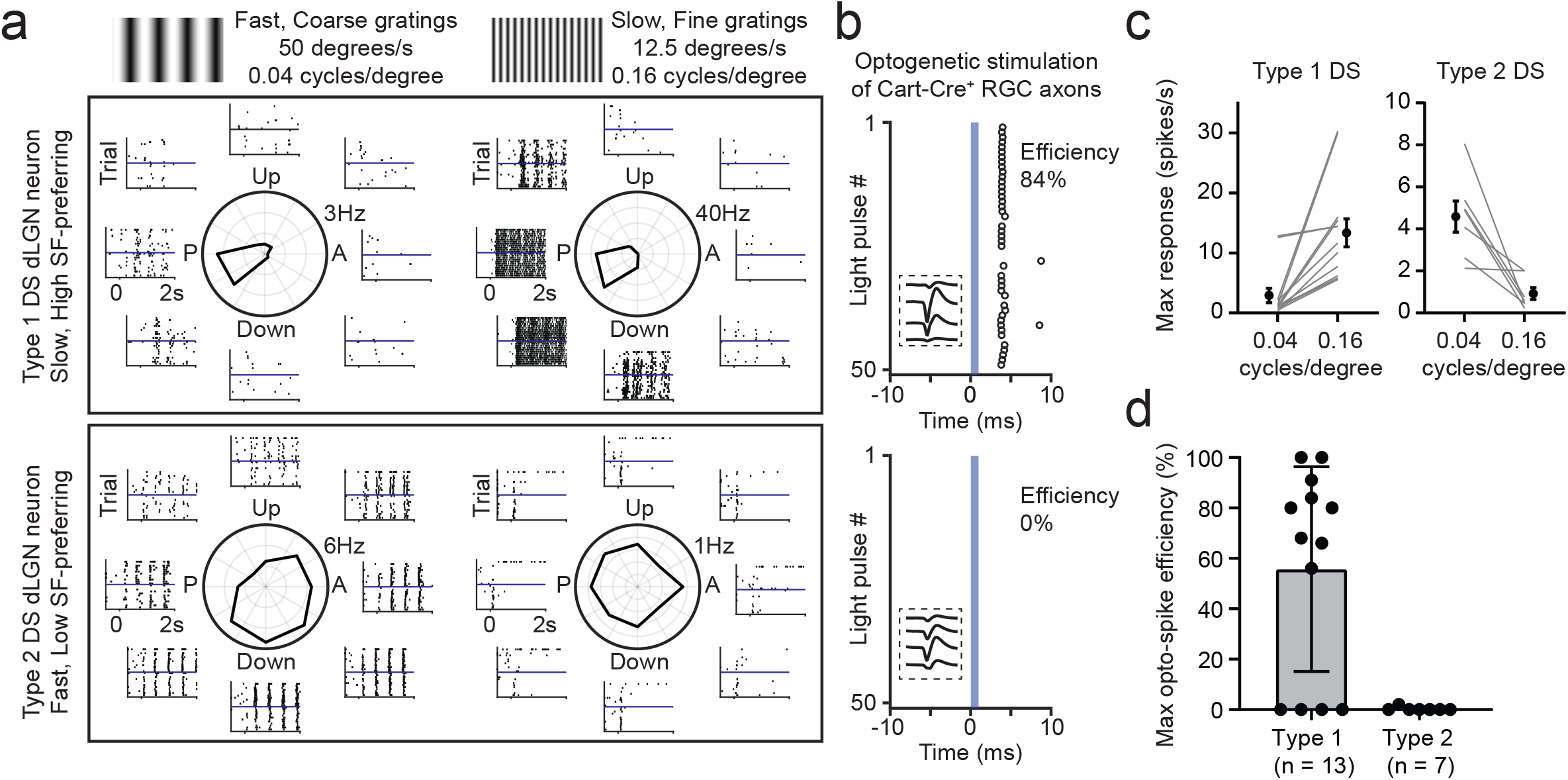
Cart-Cre^+^ inputs onto different DS types in the mouse dLGN. **a,** Example responses of DS dLGN neurons to two types of gratings. The left panels show responses to fast, coarse gratings and the right panels show responses to slow, fine gratings. The top panels are example responses of a DS dLGN neuron that prefers slow, fine gratings (type 1). The bottom panels show responses of a DS dLGN neuron that prefers fast, coarse gratings (type 2). **b,** Raster plots showing responses of the same dLGN neuron in panel (a) to 1 ms optogenetic stimulation of RGC axons. Time 0 marks the onset of optogenetic stimulation. The insets surrounded by dotted boxes are the average spike waveforms of each neuron recorded across multiple recording sites on the multielectrode array. **c,** Grouped data showing responses of type 1 (left) and type 2 (right) DS dLGN neurons to the two types of gratings. **d,** Max opto-spike efficiency of type 1 and type 2 DS dLGN neurons in response to optogenetic activation of Cart-Cre^+^ RGC inputs. Most type 1 DS neurons (9/13) receive strong input from Cart-Cre^+^ RGCs. Type 2 DS neurons were not driven by Cart-Cre^+^ inputs.

**Extended Data Figure 3.**
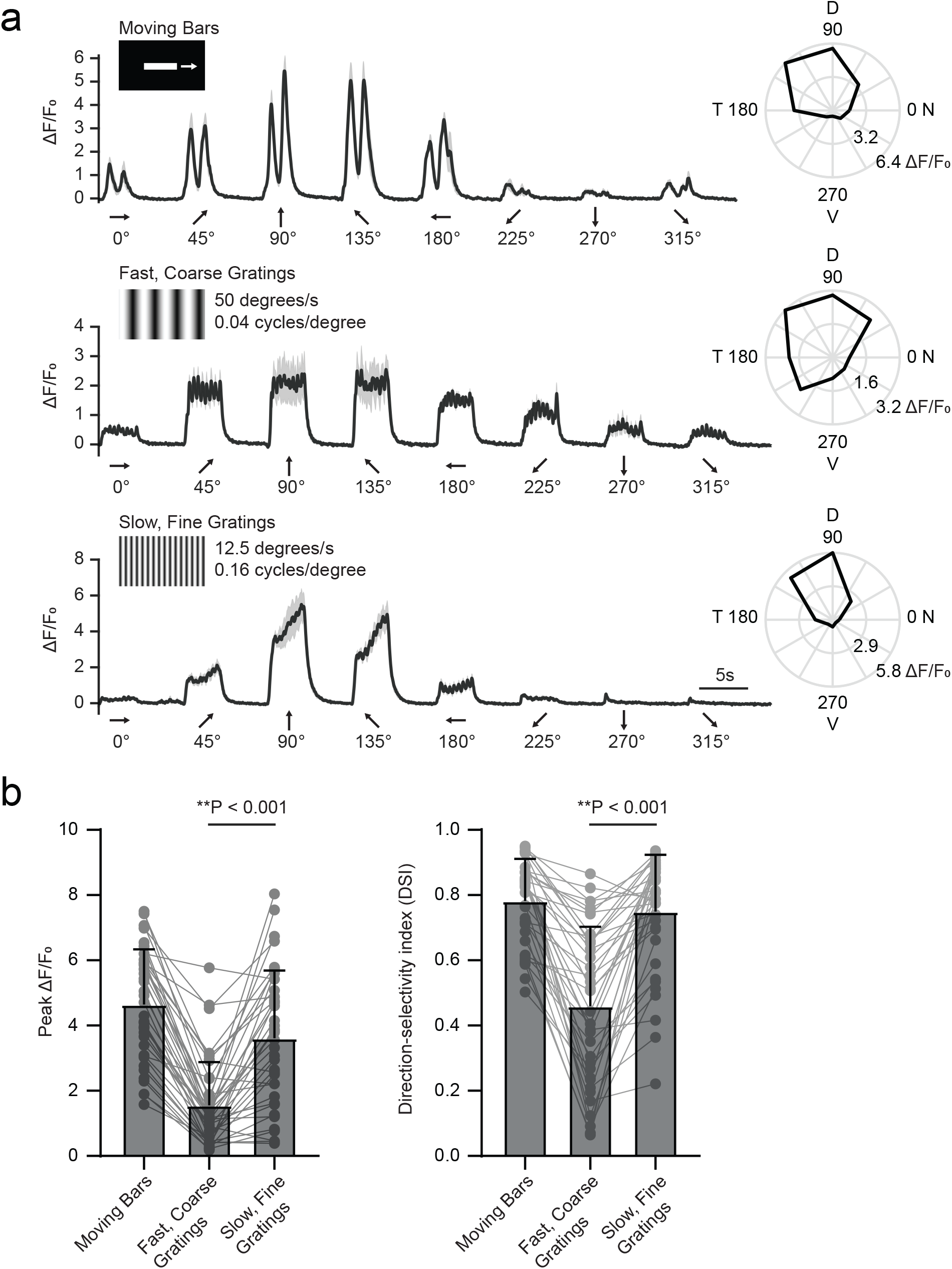
Responses of ooDSGCs to drifting gratings of different spatial frequencies. **a**, Example calcium responses of an ooDSGC labeled in Cart-Cre; Gcamp6f mice. The cell was presented with bars and drifting gratings moving in different directions. The black line represents the mean response averaged across 4 trials and the gray line represents standard error. Polar plots to the right are peak ΔF/F0 responses to stimuli moving in different directions. Note lower response amplitudes in response to faster, coarse grating stimuli. **b,** Grouped data of peak ΔF/F0 (*left*) and DSI (*right*) to different visual stimuli. ooDSGCs exhibited significantly larger responses to finer 0.16 cycles/degree gratings like type 1 DS dLGN neurons (Extended Data Fig. 2). n = 38 cells.

**Extended Data Figure 4.**
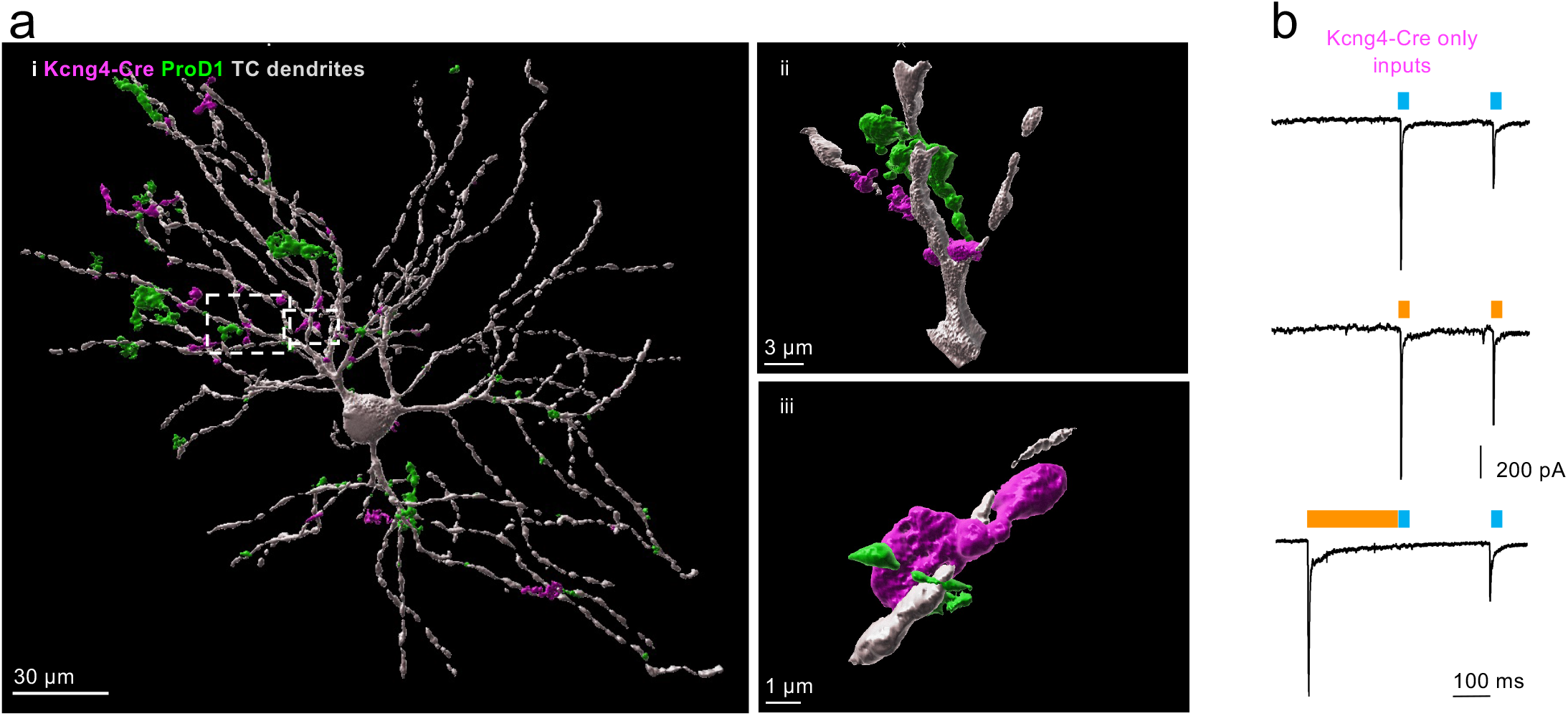
Innervation of TC neurons by Kcng4-Cre+ and ProD1+ inputs. **a**, Anatomical interaction between TC neuron dendrites and axonal terminals from Kcng4-Cre+ or ProD1+ RGCs. i) 3D reconstruction of filled TC neuron and Kcng4-Cre+ and ProD1+ terminals. Scale bar, 30 μm. Estimated surface-to-surface contact area is 647 μm^2 between TC dendrites and Kcng4-Cre+ terminals, and 872 μm^2 between TC dendrites and ProD1+ terminals. ii-iii) Enwrapping of TC dendrites by both Kcng4-Cre+ and ProD1+ terminals zoomed from dashed line boxes in (i). ii Scale bar, 3 μm; iii Scale bar, 1 μm. **b,** Functional response to Kcng4-Cre+ and ProD1+ inputs in the same dLGN neuron from panel (a). The filled neuron only receives strong inputs from Kcng4-Cre+ (812 pA), but not ProD1+ RGCs (0 pA), suggesting that anatomical interaction cannot estimate functional responses.

**Extended Data Figure 5.**
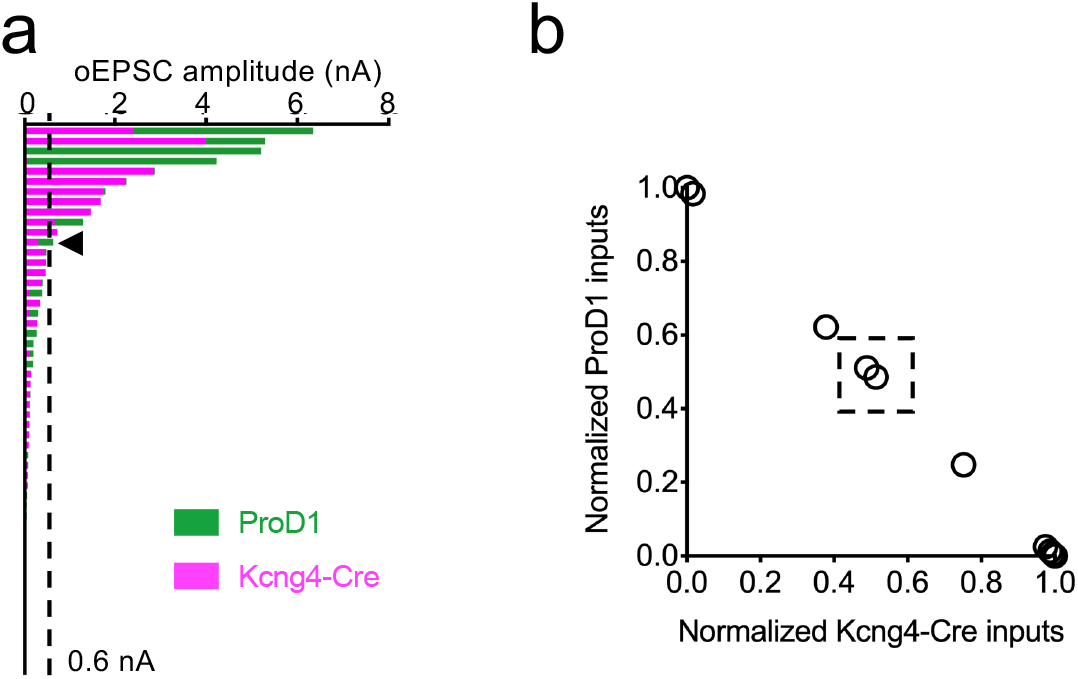
Convergence between Kcng4-Cre+ and Cart-Cre+ inputs. **a,** Distribution of oEPSC amplitude with summed responses from Kcng4-Cre+ and ProD1+ inputs. Vertical dashed line thresholds responses smaller than o.6 nA. Arrow indicates the only response with total amplitude larger than 0.6 nA but added up by two weak inputs from Kcng4-Cre+ and ProD1+ RGCs. 22% of TC neurons receive convergent inputs from these two types of RGCs regardless of their strength. **b,** Relative contribution of Kcng4-Cre+ and Cart-Cre+ inputs to gross responses with amplitude larger than 0.6 nA. Among the recordings with total amplitude greater than 0.6 nA, 83% receive dominant contribution from either Kcng4-Cre+ or Cart-Cre+ inputs.

